# Optimized data representation and convolutional neural network model for predicting tumor purity

**DOI:** 10.1101/805135

**Authors:** Gerald J. Sun, David F. Jenkins, Pablo E. Cingolani, Jonathan R. Dry, Zhongwu Lai

## Abstract

Here we present a machine learning model, Deep Purity (DePuty) that leverages convolutional neural networks to accurately predict tumor purity from next-generation sequencing data from clinical samples without matched normals. As input, our model utilizes SNP-based copy number and minor allele frequency data formulated as a scatterplot image. With a representation matching that used by expert human annotators, we best an existing algorithm using only ~100 manually curated samples. Our simple, data-efficient approach can serve as a straightforward alternative to traditional, more complex statistical methods, for building performant purity prediction models that enable downstream bioinformatic analysis of tumor variants and absolute copy number alterations relevant to cancer genomics.

## Introduction

To facilitate personalized medicine for oncology, it is critical to find novel disease-related genetic variants from patient genomic next-generation sequencing data to identify disease biomarkers, progression, or response. Hindering this effort to characterize unknown variants is an uncertainty of whether a given variant is a preexisting germline mutation or a somatic mutation arising in the tumor. To unambiguously determine germline versus somatic origin, current methodologies require a matched normal tissue sample [1–9]. However, in current clinical practice, tumor samples are collected without matched normal tissue, making it difficult to derive variant origin. Leveraging the fact that tumor samples contain a mixture of both tumor and normal cells, other methods use statistical modeling to predict the germline or somatic origin of a given variant, given the expected local copy state (i.e. tumor ploidy) and tumor/normal cell proportion (i.e. tumor purity) that best fits the observed variant allele frequency [10, 11]. These automated methods, although powerful, still require manual inspection of results, as they may yield results inconsistent with those generated by an expert human annotator. Specifically, these methods may incorrectly estimate tumor ploidy and purity, especially with noisy samples, in many cases where a human annotator might easily make a correct estimation. Lastly, only one of these methods, PureCN [10], is a fully open-source, non-commercial product available for general use.

Modern statistical machine learning methods, most notably convolutional neural networks (CNNs), have delivered revolutionary performance on unstructured data problems, such as those involving images or text [12]. Such unstructured data are often trivial for a human to interpret, but difficult for classical statistical methods to model. We therefore reasoned that for unstructured data of copy number variation and minor allele frequency, a human-suitable representation coupled with a CNN may yield performance that exceeds current statistical models. We built and trained a CNN regression model to output tumor purity based upon the very scatterplot images of input data that an expert human annotator would use for prediction. We found that with our data representation, combined with a CNN and data augmentation, we could predict purity better than an existing algorithm [10].

## Results

Using genomic data from ovarian cancer clinical FFPE tumor samples sequenced with the Foundation Medicine FoundationOne CDx panel [13], we first created a training ‘ground truth’ dataset by manually annotating 102 patient samples for purity. This was done by visualizing scatterplots of the data (Fig. 1) and comparing against expected theoretical distributions of data, given the hypothesized local copy state and observed minor allele frequencies as governed by the following equation

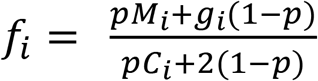

*i*: ith variant
*g*: flag for germline (1) or somatic (0)
*p*: tumor purity
*C*: local total copy number
*M*: copies of ith variant (*M* ≤ *C*)
*f*_*i*_: expected allele frequency of ith variant

**Figure 1.**
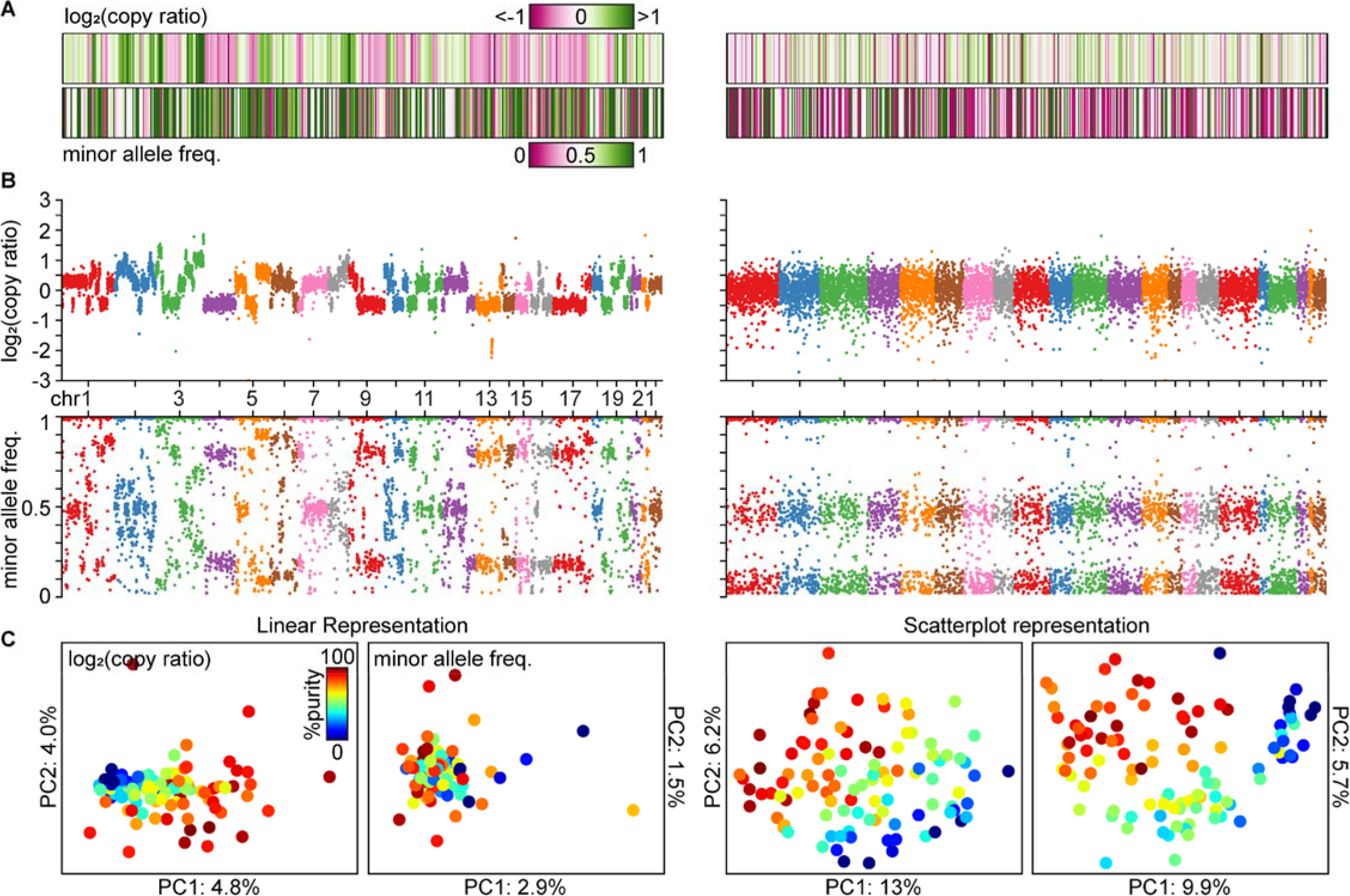
**A:** Heatmaps of raw data vectors from two ovarian cancer samples (left and right). Each vertical bar represents one real value for associated with a genomic locus. **B:** Scatterplots of data from the same two ovarian cancer samples as in A, color-coded by chromosome. **C:** Latent PCA space from the first two principal components from 102 ovarian cancer samples of either the raw data (left) or scatterplot images (right), colored by purity, separated by log_2_(copy ratio) or minor allele frequency data.

We then sought to build a model to predict purity by learning the statistics of the input data, as opposed to hand-crafting a statistical model, as with previous methods [10, 11]. Additionally, given that expert human annotators use scatterplots to visualize the data, we reasoned that the very same scatterplots would and should serve as the best data representation for our model.

With scatterplots, individual datapoints tend to cluster in space at certain allele frequencies or log copy number ratios, despite the fact that nearest neighbors in chromosome order are not necessarily similarly valued (Fig. 1). This representation therefore naturally yields a 2D histogram (Fig. 1B): datapoints in close spatial proximity cause varying degrees of spatial binning, given a particular dot size and density, such that additional datapoints in a given region yield no additional information nor appreciably change the real value of that particular ‘bin’.

In contrast, the native representation of the data—a vector of real values—is almost uninterpretable when viewed as a 1D histogram or heatmap (Fig 1A). To test this intuition and choose an optimal data representation for modeling, we generated input 1D or 2D data representations by concatenation of vectors or stacking of scatterplots, respectively. We reduced the input 1D or 2D data representations to two dimensions via principal component analysis (PCA), and found that the latter yielded a surprisingly well-structured latent space whereby tumor purity varies linearly as a combination of the principal components (Fig 1C, right). In contrast, the 1D histogram representation yielded an inferior latent space (Fig 1C, left) whereby purity did not map as cleanly.

To quantitatively assess the benefit of this data representation, we fit a simple ordinary least squares (OLS) linear regression model to the first three principal components, and computed the mean absolute error (MAE) of the OLS models’s purity predictions and our ground truth manually curated predictions for either the 1D or 2D representation. Across a 5-fold cross validation, the mean MAE of the 1D representation greatly exceeded that of the 2D representation (16.2 ± 3.3% vs 12.8 ± 1.5%; mean ± standard deviation), consistent with our qualitative assessment of superiority of the scatterplot representation.

We next optimized the scatterplot representation for our model. We systematically varied the amount of spatial binning (scatterplot image size) and the clipping threshold (scatterplot dot size) to minimize the MAE of an OLS linear regression. After optimizing our representation, we trained a simple convolutional neural network to predict tumor purity. Our trained network (5-fold cross validation) yielded 10.0 ± 1.6% MAE on the validation set, on average, across the 5 folds (Table 1). Fully-connected networks or a 1D CNN (using the native data representation vector) were unable to match the performance of a 2D CNN (14.3 ± 3.0 and 11.3 ± 2.6% MAE, respectively), despite having the same number of layers and across a wide range of trainable parameters; their performance was similar to or worse than that of our simple OLS linear regression on the first three principle components of the 2D data representation (12.8 ± 1.5% MAE). These results provide additional, quantitative evidence of the power of our chosen 2D scatterplot data representation.

**Table 1.** Mean ± standard deviation (S.D) of the mean squared error (MAE) for validation set performance of 5-fold cross validation for ovarian cancer data, across 10 runs with the same train/test split, where applicable.

| Model | MAE ± S.D. (%) |
| --- | --- |
| PureCN [10] | 8.7 |
| PCA + OLS (1D data) | 16.2 ± 3.3 |
| PCA + OLS (2D data) | 12.8 ± 1.5 |
| Fully connected NN | 14.3 ± 3.0 |
| 1D-CNN | 11.3 ± 2.6 |
| 2D-CNN | 10.0 ± 1.6 |
| 2D-CNN + augmentation | **7.7 ± 1.7** |

We next augmented our dataset by a random combination of (1) shuffling the chromosomes whilst maintaining data order within a chromosome, (2) reversing the data order of a given chromosome, (3) adding random uniform noise to the data, and (4) globally flipping the 2D scatterplot along the vertical axis, or (5) globally flipping just the minor allele frequency plot along the horizontal axis. We tested various orders of magnitude of augmentation and found that 50,000 augmented samples from our pool of 102 input samples yielded the best performance. Our final mean MAE, 7.7 ± 1.7%, was lower than that of an existing algorithm (PureCN, 8.7%).

We next examined what the model learned by computing gradient-based saliency maps for input samples [14]. We found that areas at the top edges of >2 copy segments, at the top or bottom edges of 2 copy segments, and bottom edges of 1 copy segments of the log copy number ratio plots were highlighted as being predictive for a change in purity, consistent with theory and the approach used by human annotators (Fig 2); a similar pattern consistent with theory was observed for the minor allele frequency portion of the scatterplots. Importantly, as a sanity check [15], random model weight initialization or training a model with shuffled labels yielded noisy, uninformative saliency maps (Fig 2); saliency maps were also less distinct if training the model without data augmentation (not shown). Thus through this post-hoc qualitative examination, the model learns the relevant, predictive features of tumor purity.

**Figure 2.**
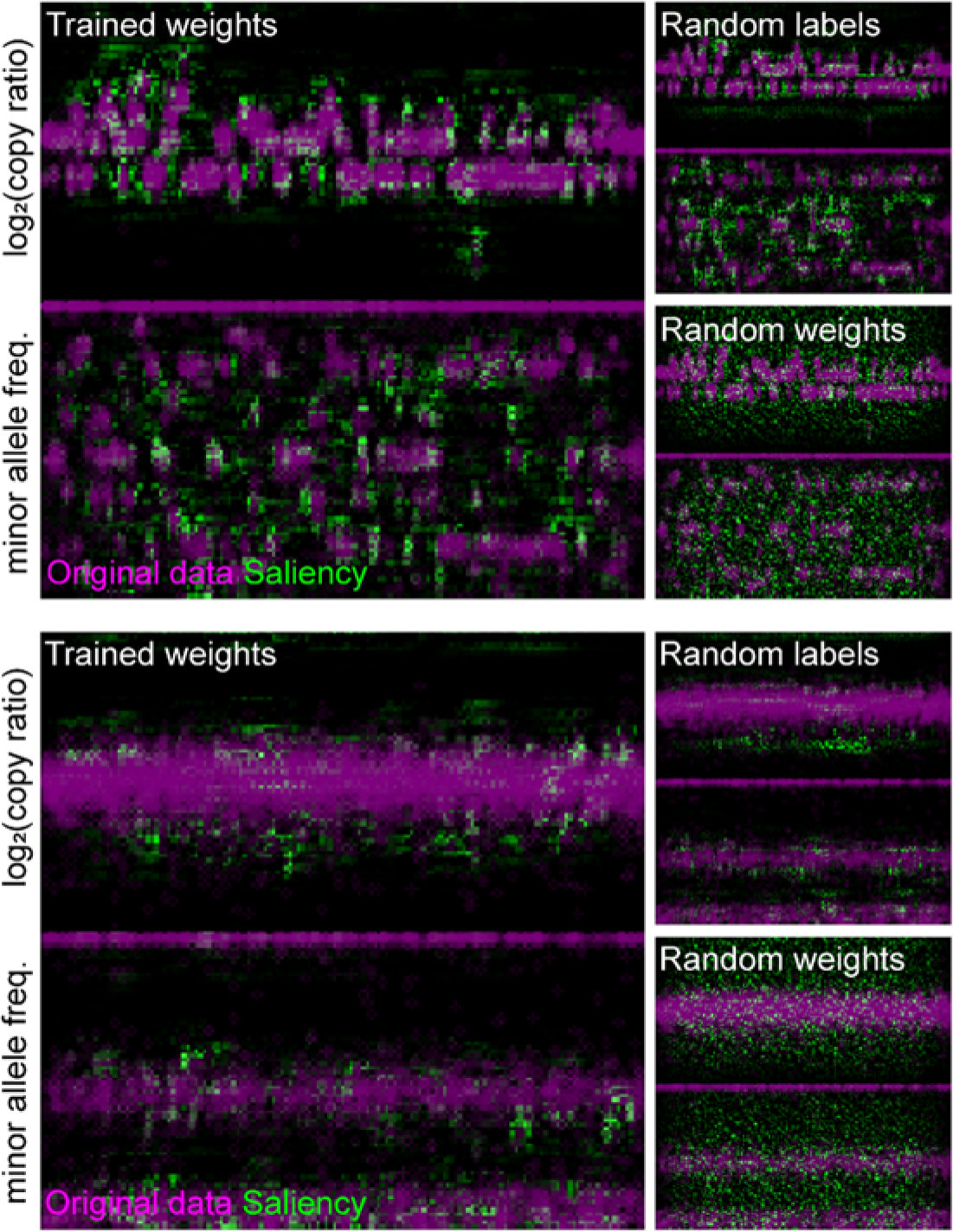
Saliency maps of the two ovarian cancer samples shown in Figure 1, as computed from the 2D CNN trained with data augmentation.

## Discussion

We found that with just ~100 training samples and data augmentation, we were able to train a performant model predicting tumor purity from gene panel sequencing of clinical tumor samples, likely due in large part to the optimal input data representation. In practice, we found that the improved tumor purity estimation (versus PureCN) increased the number of correct germline versus somatic variant determinations for known germline or somatic mutations in BRCA1/2 and TP53, respectively, in ovarian cancer (data not shown).

It is likely that a model trained solely on ovarian cancer data will not generalize well to other cancers’ genomic data. We found that this is indeed the case, and can be addressed by including other cancers’ data in the training set; as an expected byproduct, including other cancers’ data also improves model performance on the original ovarian cancer data (data not shown). Finally, our manually annotated training set represents a pseudo-ground truth, as the real ground truth tumor purity is not available. Our manual annotations do however represent a best human estimate, which is often treated as ground truth when analyzing such data as relevant to clinical trials. We purposefully did not train our model on *in vitro* tumor/normal cell admixtures, as these are not representative of data seen in the clinic. Our goal was to build and we have built a data representation and model that can robustly determine tumor purity from real-world clinical samples.

